# Modular effects of gene promoters and chromatin environments on noise in gene expression

**DOI:** 10.1101/2021.04.29.441875

**Authors:** Siqi Zhao, Zachary Pincus, Barak A Cohen

**Affiliations:** The Edison Family Center for Genome Sciences, Washington University School of Medicine in St. Louis, USA; Department of Genetics, Washington University School of Medicine in St. Louis, USA; Department of Developmental Biology, Washington University School of Medicine in St. Louis, USA

## Abstract

Genetically identical cells growing in the same environment can have large differences in gene expression. Both locally acting *cis*-regulatory sequences (CRS) and the regional properties of chromosomal environments influence the noisiness of a gene’s expression. Whether or not local CRS and regional chromosomal environments act independently on noise, or whether they interact in complex ways is unknown. To address this question, we measured the expression mean and noise of reporter genes driven by different CRS at multiple chromosomal locations. While a strong power law relationship between mean expression and noise explains ~60% of noise for diverse promoters across chromosomal locations, modeling the residual mean-independent noise suggests that chromosomal environments have strong effects on expression noise by influencing how quickly genes transition from their inactive states to their active states and that the effects of local CRS and regional chromatin on noise are largely independent. Our results support a modular genome in which regional chromatin modifies the inherent relationship between the mean and noise of expression regardless of the identity of the promoter sequence.

## Introduction

Genetically identical cells growing in the same environment can display large differences in the mRNA levels of expressed genes(Elowitz et al., 2002; Raj et al., 2006; Zenklusen et al., 2008). This cell-to-cell variance in mRNA abundance among isogenic cells is called noise in gene expression and drives important aspects of biology. Cell fate decisions for olfactory sensory neurons(Chess et al., 1994), retinal pigment cells(Wernet et al., 2006), intestinal crypt cells(Tóth et al., 2017), neural precursors in the spinal cord(Dasen et al., 2003, 2005), and hematopoiesis(Chang et al., 2008) are governed by the noisy expression of lineage-determining transcription factors. Random fluctuations in gene expression also affect how individual melanoma cells respond to chemotherapy(Shaffer et al., 2017). Single-cell transcriptome profiles from scRNA-seq suggest that expression noise is universal for many cell types(Cembrowski and Menon, 2018; Grün et al., 2015; Villani et al., 2017) and likely accounts for different cell states within the same cell type(Adler et al., 2019). Noise in gene expression is thus an important component of both development and disease.

Noise in gene expression results from random fluctuations in the molecular processes that control transcription. Single-cell studies show that transcription is not continuous, but instead occurs in discrete bursts of activity(Lenstra et al., 2016; Zoller et al., 2018). A gene’s promoter sequences(Sharon et al., 2014; Tunnacliffe et al., 2018), its epigenetic modifications(Nicolas et al., 2018), or its patterns of DNA looping(Fukaya et al., 2016) can all influence the burst size and frequency of genes. For different genes, the burst frequency can range from just a few minutes to tens of hours(Lenstra et al., 2016; Rodriguez et al., 2018). Consequently, cells in the midst of bursting will have higher mRNA counts than cells waiting for their next burst of expression. The bursty dynamics of mammalian transcription is an important source of expression noise.

Two types of regulatory information can influence expression noise: locally acting CRS, such as promoters and proximal enhancers, and more distally acting factors, such as long-range enhancers and sequences that control the regional chromatin state. In this study, we define the regional “chromosomal environment” as the combination of a gene’s distally acting enhancers and its broader chromatin landscape. Classic studies of Position Effect Variegation (PEV) show that different chromosomal environments can have large effects on noise(Wallrath and Elgin, 1995). More recent work has shown how features associated with local *cis*-regulatory sequences (CRS), such as the TATA-box(Hornung et al., 2012), the number of transcription factor binding sites(Soltani et al., 2015), the local nucleosome occupancy(Dey et al., 2015), and local epigenetic modifications(Nicolas et al., 2018) can all affect expression noise. Two studies used randomly integrated reporter genes to show that different chromatin environments can affect expression noise (Dar et al., 2012; Dey et al., 2015). However, how the effects of regional chromosomal environments combine with the effects of local regulatory features to control noise remains an open question.

Local and regional *cis*-regulatory information may act independently to control noise in expression, or there may be complex interactions between local CRS and their regional chromosomal environments. If these two sources of *cis*-regulatory information act independently on noise, then the effects of chromosomal environments should be the same for different classes of local CRS. Alternatively, a gene’s noise may depend on complex interactions between local CRS and their chromosomal environments: certain locally acting *cis*-elements may only increase noise in certain chromosomal environments. We previously showed that local and regional *cis*-regulatory information act independently to control mean levels of expression(Maricque et al., 2018). In this study we sought to determine whether the modularity between local and regional *cis*-regulatory information extends to the control of noise in expression.

To address this question, we constructed a series of reporter genes driven by different promoters and integrated them into several defined genomic locations in the K562 cell line. We measured both the mean and noise of each reporter gene at each genomic location with single molecule Fluorescence in situ Hybridization (smFISH). Our results show that across all reporters at all locations, ~60% of expression noise depends only on the mean level of the reporter gene and is captured by a power law. The residual mean-independent noise is mainly associated with the epigenetic signature of the regional chromosomal environment. Given the same mean level of expression, more active chromatin environments generally result in less expression noise than more repressive environments. By comparing diverse reporter genes integrated at the same chromosomal locations we found that the effects of different chromosomal environments on expression noise were independent of the specific promoter. Taken together our results support the notion of a modular genome in which local and regional *cis*-regulatory information act independently to control both mean levels of expression and noise in expression.

## Results

### An experimental system to quantify noise at diverse chromosomal locations

We constructed a system to separate the contributions of locally acting CRS and regional chromosomal environments to expression noise. We used a set of previously generated landing pad cell lines that contain reporter genes integrated in defined chromosomal locations across the K562 genome (Maricque et al., 2018). Each landing pad cell line has an identical reporter gene cassette in a different mapped genomic location (**Fig. 1 A, B**). In this study, we used landing pads integrated at twenty-two different genomic locations, with different epigenetic marks (**Fig.S1**).

**Fig 1.**
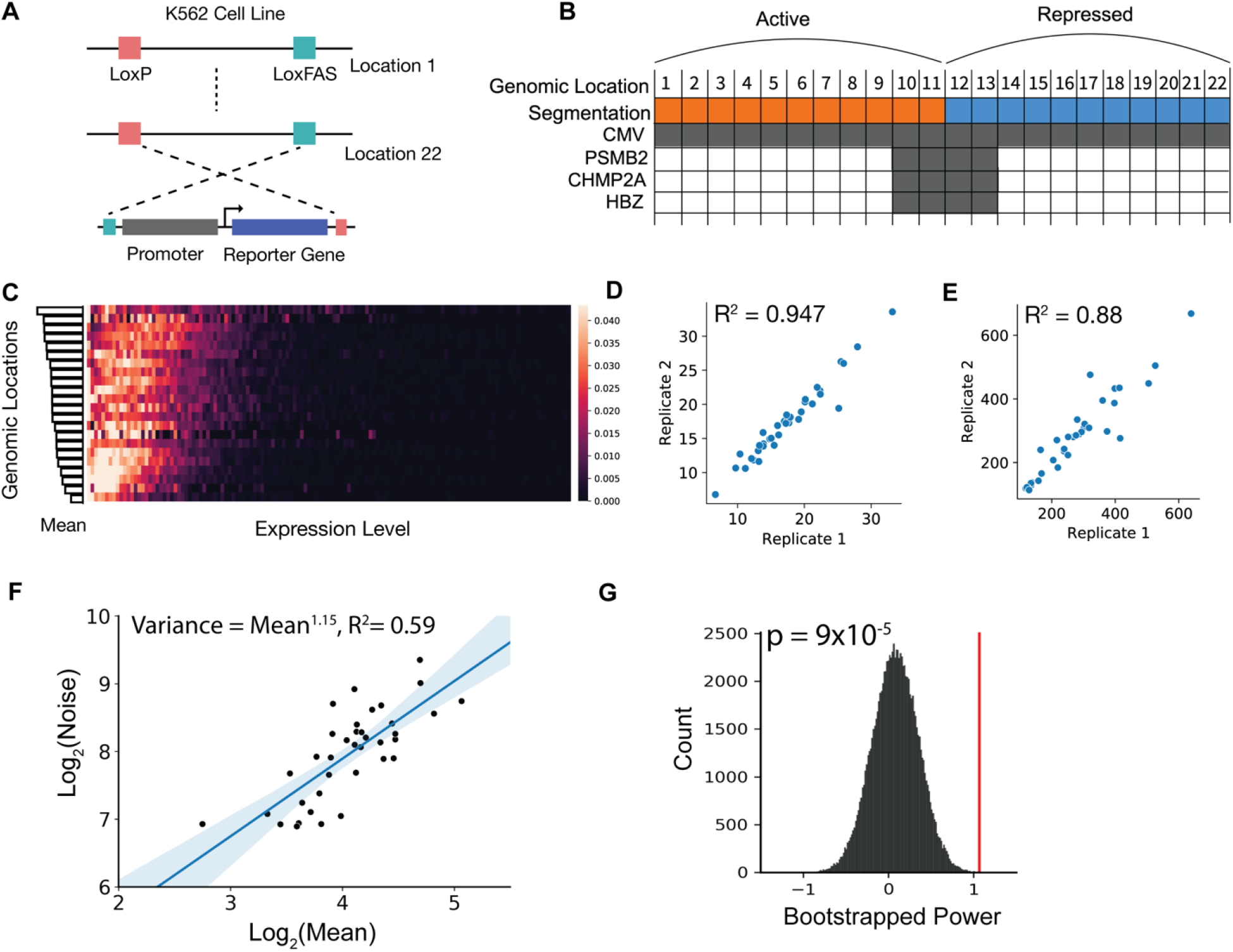
Experimental design for studying the cell-to-cell variability associated with different genomic locations. **(A)** Strategy for integrating the same reporter cassette into mapped genomic locations. In each K562 cell line, a construct containing a pair of LoxP and LoxFAS sites is integrated and mapped. The sequence between the pair of lox sites contains a reporter cassette of eGFP driven by a CMV promoter. Three other reporter cassettes containing PSMB2, HBZ, and CHMP2A promoters replace the CMV promoter through cre recombination. **(B)** Schematics for selecting K562 cell lines. 8 active locations and 14 repressed locations were selected based on the ChromHMM+Segway combine segmentation (see Methods). **(C)** Heatmap of single-cell mRNA expression of 22 cell lines with CMV promoter, ordered from highest mean expression level to lowest. Each position in the matrix represents the mRNA counts of a single cell, and the color represents the percentage of cells. **(D, E)** Scatter plots showing the reproducibility derived from two replicates for mean (**D**) and noise (**E**) of the measured single-cell expression distributions. **(F)** Linear regression of mean against noise in the log scale revealed a power-law dependence. Mean expression level explains about 59% of the observed noise, with the power of 1.15. **(G)** Histogram of the bootstrap of the power of the power-law relationship between mean and noise. The red line shows the fitted power from experimental data.

Each landing pad carries a common reporter gene consisting of a locally acting *cis*-regulatory sequence (CRS) driving the expression of Enhanced Green Fluorescent Protein (eGFP) and flanked by a pair of asymmetric *Lox* sites allowing us to easily exchange the reporter gene cassette. (**Fig. 1A**). For all twenty-two locations, we integrated an eGFP reporter driven by the cytomegalovirus (CMV) promoter. To explore the dependence of noise on local CRS, we integrated three other reporter cassettes with tdTomato reporter gene driven by PSMP2, HBZ, and CHMP2A promoters at four locations (**Fig. 1B**). To simplify the system, we intentionally chose not to include an intron in the reporter cassette to avoid potential confounding effects of different genomic locations on the efficiency of splicing(Mikl et al., 2019). With this system, we can systematically probe how different chromosomal environments affect the activities of different local CRS.

One of the challenges of studying the effect of transcriptional regulation is to exclude the confounding effects of post-transcriptional regulation. Hence, for each reporter gene at each chromosomal location we performed clampFISH, a smFISH method, to measure the absolute mRNA counts in individual cells (Rouhanifard et al., 2018). To accurately quantify both expression mean and noise, we quantified mRNA molecules from more than 8×10^5^ cells and obtained more than 1000 single-cell mRNA measurements per cell line per replicate. The experimental measurements are highly reproducible for both mean and noise at each genomic location (R^2^ = 0.947 for mean, R^2^ = 0.88 for noise, **Fig.1 D, E**). The absolute measurements of mRNA molecules using smFISH allowed us to use computational modeling to infer transcriptional dynamics.

### The mean and noise of gene expression are linked by a power-law relationship

We observe dramatic changes in the mean and noise of expression that depend on genomic locations. We observed a 6-fold difference in mRNA mean and a 7-fold difference in noise among the genomic locations we sampled (**Fig. 1C**). This dynamic range of mean and noise is consistent with the range observed in a previous study where the investigators measured mRNA in cell lines with random integrations in a mammalian cell line (Dey et al., 2015). Thus, our data is consistent with both classic work on PEV and more recent single-cell studies, which show that changing the genomic location of a gene has large effects on its expression mean and noise.

We first established the general relationship between the mean and noise of expression in our data. In all stochastic processes there is a strong dependency of noise on the mean output levels. Because changing the genomic location of a gene affects its mean expression, its noise will also necessarily change due to the dependency of the noise on the mean. Therefore, it is important to establish the general relationship between expression mean and noise in our experiments so that we can decompose the effects of genomic location on noise into effects that are due solely to changes in mean expression levels and those that are independent of mean level changes.

To characterize the general relationship between the mean and noise in our data we performed a log-log regression of expression noise on mean expression levels across all CRS at all genomic locations. Consistent with previous findings, this analysis revealed a power law relating expression noise to mean expression levels. Differing from studies using fluorescence-based single-cell quantification of genome-wide protein abundance, which report a degree of 1.96, we observed a smaller degree of 1.15 (noise = mean^1.15^, R^2^ = 0.6) (**Fig. 1F**) (Das et al., 2017; Dey et al., 2015; Vallania et al., 2014). To determine the significance of the fitted degree for the power law relation between expression mean and noise, we performed 10^5^ bootstrap simulations and obtained a p-value of 9*10^−5^ for the fit to the actual data (**Fig. 1G**). We interpret the higher degree for the power law relationship for the protein distribution to represent additional noise introduced by post-transcriptional steps in gene expression (Paulsson, 2005). In our experiments, 60% of the noise in mRNA levels is explained solely by the effects of different chromosomal locations on mean levels. The same power law relation holds regardless of the local CRS identity: there are no promoter specific mean-noise relationships in our data. When we plot the mean-noise relationship for different promoters alone, the power law relation remains largely the same (**Fig. S2A-D**). This result suggests that a general aspect of the transcriptional mechanism accounts for a significant fraction of observed noise, regardless of the specific sequence context.

The power law’s degree of ~1.15 reflects the bursty nature of mammalian gene expression. If the reporter genes were constitutively producing mRNA, then we would expect Poisson dynamics where the degree for scaling is 1.0 (Bar-Even et al., 2006; Paulsson, 2005). The non-Poisson relationship in our data suggests the existence of “ON” and “OFF” states for transcription activation (Raj et al., 2006), which agrees with the bursty dynamics observed in mammalian cell systems and *Drosophila melanogaster (Fukaya et al., 2016; Zoller et al., 2018)*. Consistent with the existence of an OFF state, for each reporter gene at each location there was always a subpopulation of cells with no mRNA (**Fig. S2E**). This suggests that in some cells, transcripts are degraded completely before the reporter gene transitions to the ON state. To confirm that this observation is not due to false negative labeling of mRNAs, we performed smFISH on the introns of the constitutively expressed gene *Actb*, which encodes the beta actin protein. For *Actb*, less than 1% of the cells have no labeled transcript (**Fig. S2E**), which suggests that the larger fraction of cells with no reporter gene mRNAs are not false negatives resulting from experimental artifacts such as poor fixation or permeabilization. Both the observation of a non-Poisson dynamics and the existence of cells with no mRNA present suggests that there are regulated, slow steps in transcription that generate noise in gene expression. We sought to explore the mechanism behind the difference in transcriptional dynamics at different genomic locations.

### Mean Independent Noise (MIN) describes expression noise without the mean effect

The power law relation shows that mean expression levels explain 60% of expression noise. We define the residual 40% of expression noise that is not explained by mean levels as Mean Independent Noise (MIN). In all subsequent analyses we use MIN as the metric of noise because it shows no dependence on mean levels (**Fig. 2A**), while other commonly used metrics of noise, such as the Fano Factor (σ^2^/μ) or CV^2^ (σ^2^/μ^2^), still show some dependence on mean levels (**Fig. S3 A, B**). Because MIN is based on the power law fit to our data (MIN ~ σ^2^/μ^1.15^) it more naturally expresses the relationship between mean expression levels and noise. In all subsequent analyses we use MIN as a measure of the effects of chromosomal environments on expression noise.

**Fig 2.**
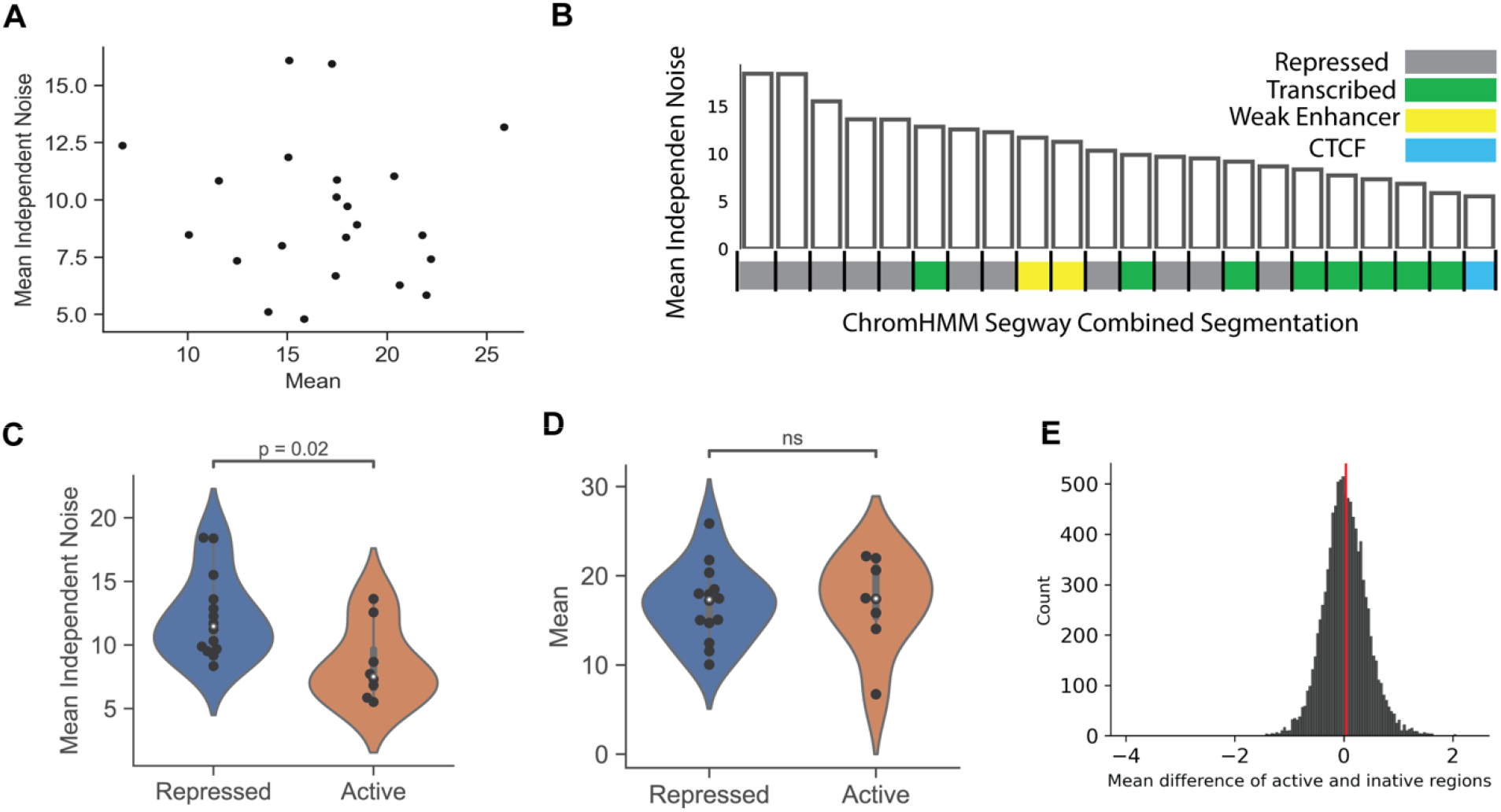
Active genomic locations have lower expression noise. **(A)** Expression mean and MIN are plotted for each genomic location expressing eGFP driven by the CMV promoter. (**B**) Residual noises from the power-law fitting are plotted with the order from highest residual to the lowest. Each bar corresponds with the combined ChromHMM+Segway segmentation annotations. **(C)** Violin plot for MIN at active and repressed locations. **(D)** Violin plot for mean at repressed and active locations. **(E)** Histogram of trip mean expression differences from Ahktar et al (2013) for 10^5^ times using the number of observations from our experimental data (n=22), red vertical line indicates the actual mean difference observed from our data.

### Chromatin states explain Mean Independent Noise at different chromosomal locations

What mechanisms control MIN? We hypothesized that regional chromatin environments have direct effects on MIN. Before analyzing whether regional chromatin environments affect MIN, we addressed whether the reporter gene causes gross alterations in the chromatin environment. We performed 4C (Zhao et al., 2006) on two cell lines containing reporter genes and found that the contact maps agree with the contact frequencies derived from the Hi-C data on cells without reporter genes (**Fig. S3 E,F**) (ENCODE Project Consortium, 2012; Rao et al., 2014). We then asked whether the different epigenetic properties of genomic locations explain the differences in expression mean and MIN at different genomic locations (ENCODE Project Consortium, 2012). We first examined individual epigenetic marks and found some active chromatin marks are enriched in active regions 5kb around the insertion site (**Fig. S4 A-F**). However, due to the sparseness of the peaks of histone modifications within 500 bp around our insertion sites, we decided to look at the aggregated epigenetic information (**Fig. S4G**). We then plotted the MIN from cell lines with CMV promoters against the genomic annotations of the 100 bp flanking the reporter genes. Genomic locations labeled as Transcribed by the chromHMM and Segway (Hoffman et al., 2012, 2013) combined annotations have lower MIN than regions labeled as Repressed (**Fig. 2B,C**). We observed similar results using either the Fano Factor or CV^2^ as the noise metric (**Fig. S3C, D**). We hence classified our genomic locations as active (Transcribed, Weak Enhancer, and CTCF site based on ChromHMM segmentations) and repressed (Ernst and Kellis, 2012; Hoffman et al., 2012, 2013), which combine diverse epigenetic regulatory information into annotations of chromosomal locations (**Supp. Table 1**).

Interestingly, we did not find strong differences in mean expression levels between active and repressed locations (**Fig. 2D**). Previous studies of the genomic location effect on expression mean have shown that reporters integrated into the active locations have statistically higher mean expression (Akhtar et al., 2013). However, those statistically higher expression mean levels are often driven by a small number of genomic locations. To illustrate this fact, we used a previously published dataset containing the mean measurement of a reporter gene integrated in thousands of genomic locations. We then subsampled the data and compared the z-score of the expression mean level at active and initiative locations. We found that there are no significant differences in mean expression levels when we randomly sample 22 locations from the whole dataset (**Fig. 2E**). Taken together, our results suggest that the epigenetic differences between chromosomal locations have larger effects on the noise of gene expression than on mean levels of expression.

### Computational model reveals the dynamics that changes cell-to-cell variability at different genomic locations

To explore the underlying mechanisms that explain the observed difference in MIN at different genomic locations, we studied how different genomic locations control the bursting dynamics of expression. Recent studies suggest that the control of transcriptional bursts directly affects expression noise (Rodriguez et al., 2018). Theoretical and experimental works show that the bursty dynamics of single-cell gene expression can be described by the two-state ON/OFF model (**Fig. 3A**) (Bar-Even et al., 2006; Paulsson, 2005; Raj et al., 2006). At the same time, this framework has successfully been used to connect single-cell variability with transcriptional dynamics experimentally (Das et al., 2017; Dey et al., 2015; Lammers et al., 2020; Nicolas et al., 2018; Raj et al., 2006). The ON/OFF model abstracts gene expression into four macroscopic processes, each with a corresponding rate constant. K_on_ and K_off_ describe the transition of chromatin between the ON and OFF states, K_m_ describes the rate of mRNA production (which only occurs when the chromatin is in the ON state), and K_d_ describes the rate of RNA degradation. These rate constants set the burst size (K_m_/K_off_) and burst frequency (K_on_/K_off_) of a gene (Dar et al., 2012, 2016).

**Fig 3.**
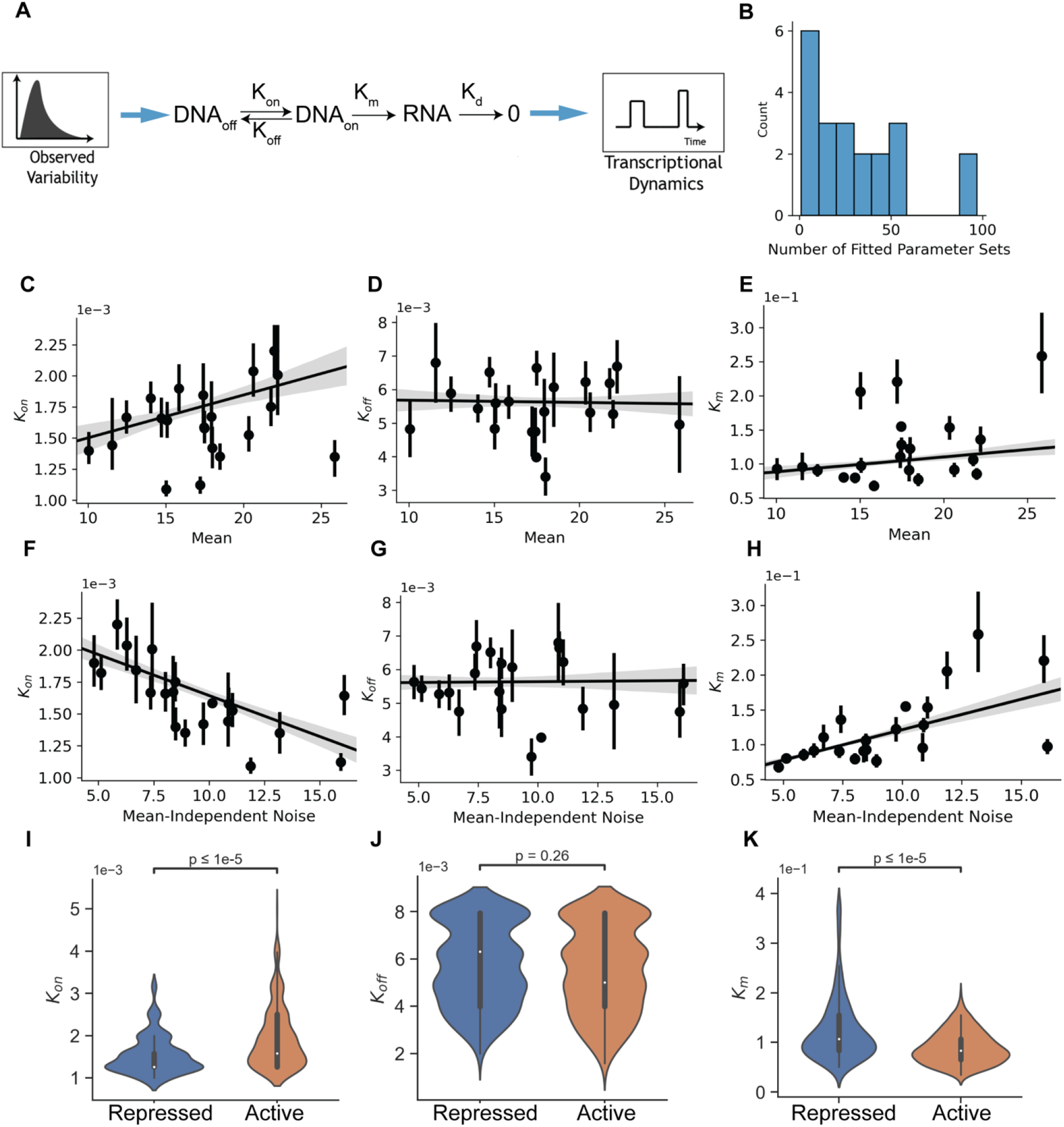
The ON/OFF model reveals the dynamics that drive different MIN at active and repressed locations. **(A)** Schematics for stochastic simulation for the two-state ON/OFF model. **(B)** Number of fitted sets of parameters for each experimentally measured distribution (n = 34, median = 23). **(C–E)** Scatter plots of mean against K_on_ (C) (R^2^ = 0.03, p = 1.7×10^−6^), K_off_ (D) (R^2^ = 0.0001, p = 0.73), and K_m_ (E) (R^2^ = 0.02, p = 0.00015),). Error bar indicates all fitted values from the stochastic simulation. Shadow shows the 95% C.I. of the linear fit. **(F–H)** Scatter plots of MIN against K_on_ (F) (R^2^ = 0.1, p = 5.2×10^−17^), K_off_ (G) (R^2^ < 1×10^−5^, p = 0.8), and K_m_ (H) (R^2^ = 0.3, p = 3.38×10^−52^). Error bar indicates all fitted values from the stochastic simulation. Shadow shows the 95% C.I. of the linear fit. **(I-K)** Violin plots of all fitted K_on_ (I), K_off_ (J), and K_m_ (K) at active and repressed locations.

We employed an exhaustive fitting strategy for the ON/OFF model to identify parameter sets that might explain the differences in MIN between different genomic locations. We found a median of 23 sets of parameters that fit each of our experimental distributions and those sets of parameters cluster distinctively based on different genomic locations (**fig. 3B**). Compared to the thousands of sets of parameters obtained from fitting those parameters with protein distribution data, our fitted result is 100-fold less degenerative, this allows us to infer the differences in dynamics at different genomic locations.

We first examined the general trend for how different parameters in the ON/OFF model correlate with mean expression levels. We found that only the transition to the ON state is positively correlated with the increase of expression mean (**Fig. 3C**); there is no change for the rate for transitioning to the OFF state (**Fig. 3D**). This suggests that the genomic location effect on mean is mainly associated with the opening of the chromatin, and there is a constant process controlling the transition to the OFF state. This result agrees with the recent study showing that the inactivation of chromatin is a constant, and active process (Falk et al., 2019). We also found a weak correlation between expression mean and K_m_, the rate of transcription (**Fig. 2E)**. Overall, the computational modeling agrees with experimental observations of how mammalian chromatin environments affect the mean of gene expression.

We next asked how the different parameters of the ON/OFF model correlate with MIN. Surprisingly, we found that the transition to the ON state is negatively correlated with MIN (**Fig. 2F**). This suggests that the faster the transition to the ON state, the lower the MIN. Thus, K_on_ appears to have opposing effects on mean and MIN, which suggests that expression mean and noise can be orthogonally controlled. We found that K_m_ is also positively correlated with MIN (**Fig. 2H**), suggesting that the rate of transcription increases both the mean and MIN of expression.

Overall, the ON/OFF model revealed that chromosomal environments mainly affect K_m_, the macroscopic parameter describing the rate of transcription, and K_on_, the parameter describing the rate of activation. However, we discovered that MIN decreases as K_on_ increases, suggesting faster activation is a potential mechanism for suppressing expression noise. Moreover, the orthogonal effect of K_on_ on expression mean and MIN suggests that mean and MIN can be decoupled by tuning different macroscopic steps of gene expression.

### Active chromosomal environment produces less expression noise by creating frequent but small transcriptional bursts

We then asked if the ON/OFF model explains the difference in MIN at active and repressed genomic locations. We found that reporter genes driven by the CMV promoter tend to have faster K_on_ when integrated in active locations compared to repressed locations, while K_off_ is not distinguishable for the transcribed and repressed locations (**Fig. 3 I, J**). This suggests that at active locations, reporter genes transition faster to the ON state than at repressed locations. However, repressed locations have slightly higher transcription rates (K_m_) compared to active locations (**Fig. 3K**). To achieve the same mean level at a repressed location with slower ON rate, the transcription rate must be higher to compensate for the lower activity of the chromatin transition.

We then estimated the transcriptional burst size and frequency from the fitted parameters (**Fig. 4C)**. Intuitively, we found positive correlations between burst size and frequency with expression mean (**Fig. 4A, B**), increasing either burst frequency or burst size can lead to the increase of mean expression, and different chromatin environments can modulate both burst size and burst frequency. Interestingly, while burst size still is positively correlated with MIN, burst frequency is negatively correlated with MIN (**Fig. 4D, E**). We also found that active locations have higher burst frequencies and lower burst sizes, while repressed locations have lower burst frequencies and higher burst sizes (**Fig. 4F, G**). This suggests that faster burst dynamics reduce MIN without reducing expression mean level. This finding provides a potential mechanism for reducing expression noise without lowering the mean expression through controlling the transcriptional dynamics.

**Fig 4.**
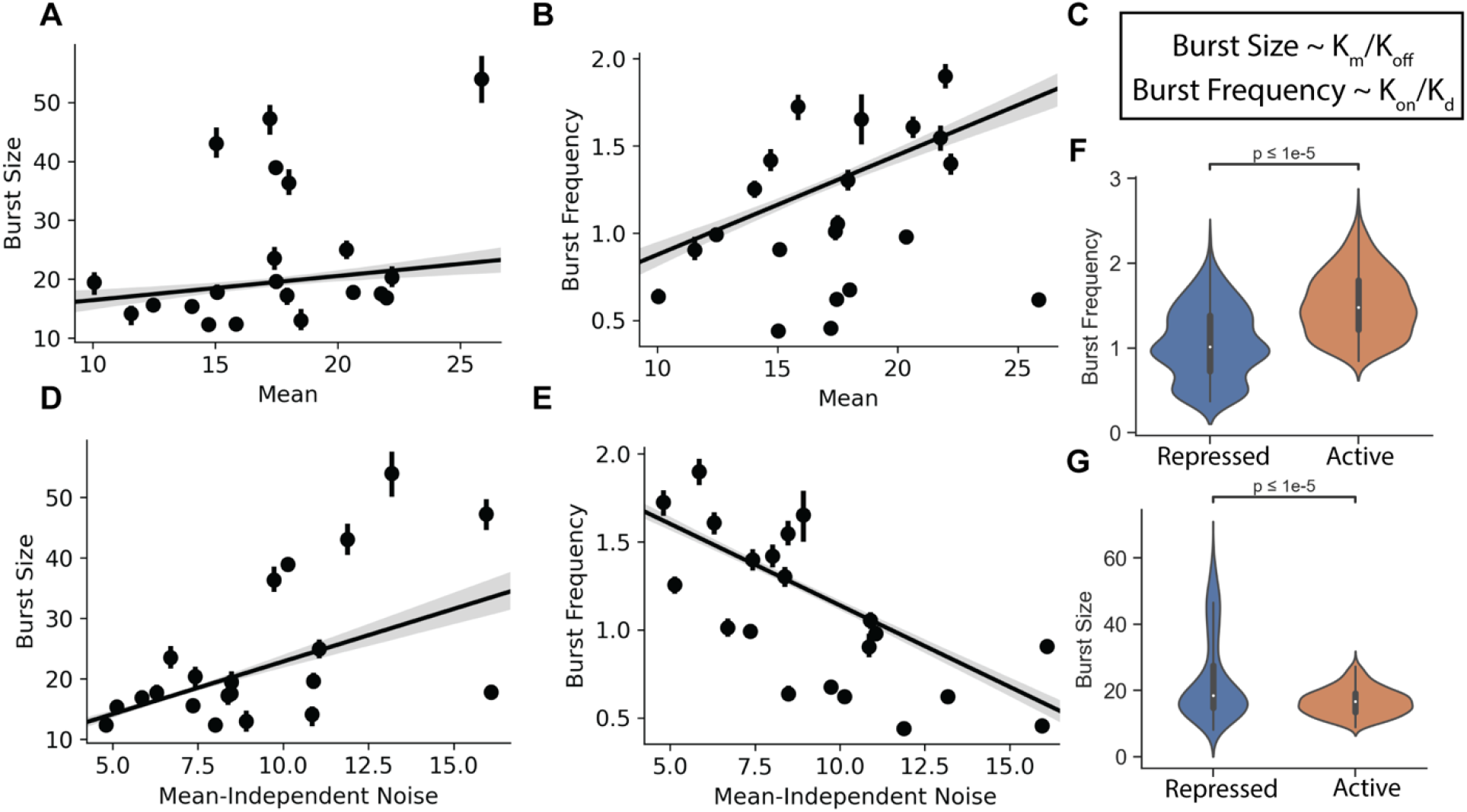
Active transcriptional locations have higher burst frequency but lower burst size. **(A, B)** Scatter plots of mean against burst size (R^2^ = 0.17, p = 0.0001) and burst frequency (R^2^ = 0.19, p = 5×10^−32^). Error bar indicates all fitted values from the stochastic simulation. Shadow shows the 95% C.I. of the linear fit. **(C)** Derivation of average burst size and burst frequency based on the ON/OFF model. **(D, E)** Scatter plots of MIN against burst size (R^2^ = 0.34, p = 7×10^−^61) and burst frequency (R^2^ = 0.43, p = 2×10^−81^). Error bar indicates all fitted values from the stochastic simulation. Shadow shows the 95% C.I. of the linear fit. **(F)** Violin plot for burst frequency at active and repressed locations. **(G)** Violin plot for burst size at active and repressed locations.

Overall, we found that expression mean and MIN are decoupled through differential control of different steps of transcription. We found that active genomic locations have MIN through faster burst dynamics, but slightly smaller burst sizes compared to repressed locations. Moreover, the faster dynamics is due to faster transitions to the ON state, and not to slower transitions to the OFF state. These results suggest that expression mean and MIN can be decoupled through increasing the ON transition, but not by increasing the rate of transcription.

### Independence of the effects of genome location and local CRS on noise in gene expression

To determine whether the effect of chromatin environments on MIN depends on the specific local CRSs, we analyzed the expression distribution of reporter genes with three other promoters (CHMP2A, PSMB2, and HBZ) at the same two active and two repressed genomic locations. Interestingly, the resulting mean-noise relationship still falls on the same power-law relationship (**Fig. S2A**). While the promoter identity determines the mean level of expression all four locations (**Fig. 5A**), we found that chromatin environments have a strong effect on MIN, regardless of the specific promoter identity (**Fig. 5B**). To quantify the contribution of local CRSs and chromatin environment on MIN, we performed a two-way ANOVA, and found that genomic location explains ~51% of the variance of the MIN (p = 0.039), while promoter identity is not a significant predictor of the observed variance of the MIN. In contrast, promoter identity explains ~65% of the variance of the mean (p = 0.0012), and the genomic location explains ~20% of variance of the mean (p = 0.046) (**Fig. 5E**). The ANOVA analysis affirms the observation that the regional chromatin environment of a gene has a strong effect on the MIN.

**Fig 5.**
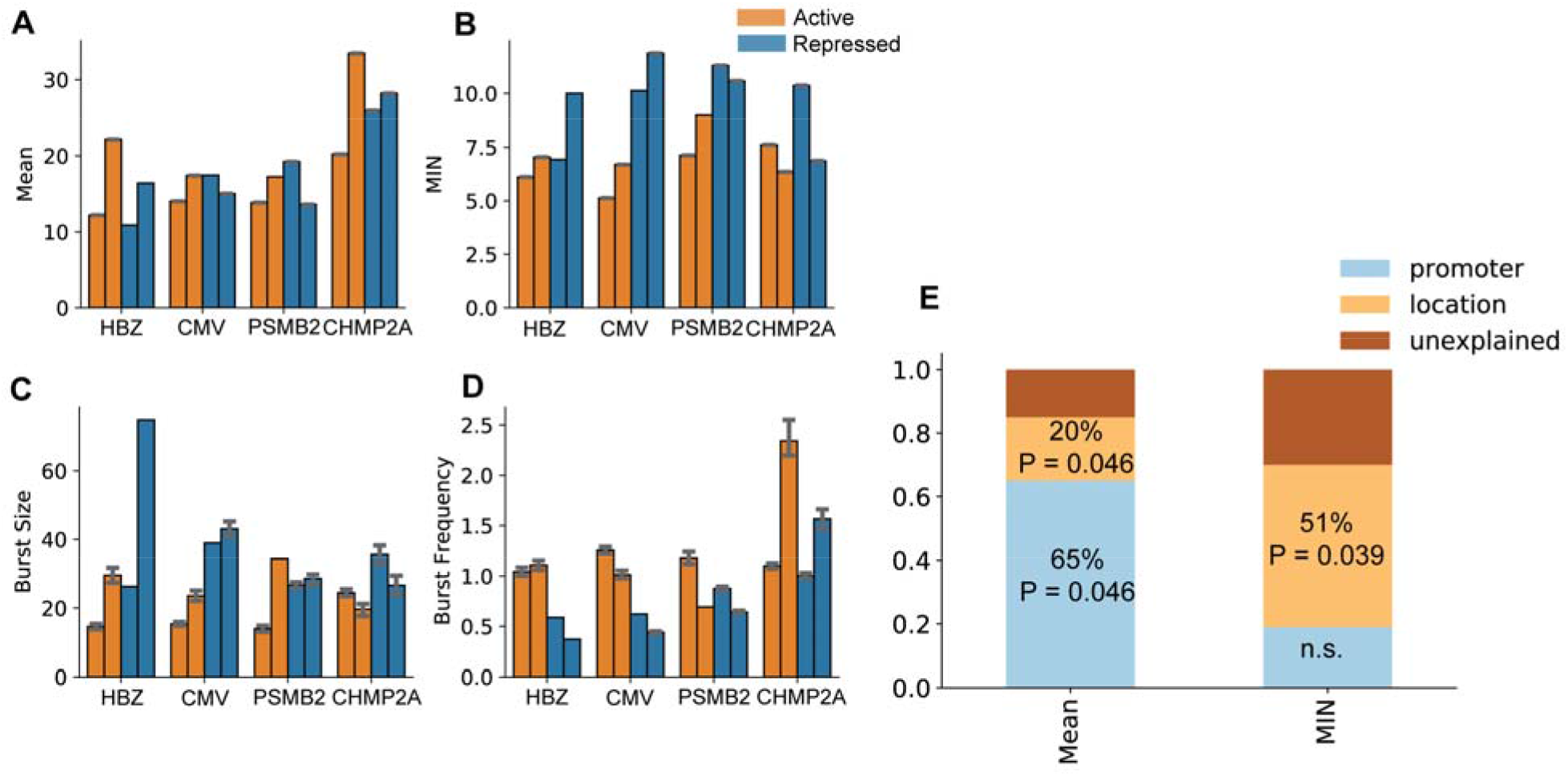
Active transcriptional locations have higher burst frequency but lower burst size. **(A, B)** Bar plots showing the mean and MIN of different promoters at two active and two initiative genomic locations. **(C, D)** Bar plots showing the burst size (C) and burst frequency (D) of different promoters at two active and two initiative genomic locations. (**E**) Two-way ANOVA analysis for promoter identity and genomic location contribution to expression mean and MIN.

Agreeing with the data from CMV promoters, reporter genes with the other promoters integrated in the active locations also have smaller burst sizes and higher burst frequencies compared to those integrated in the repressed locations (**Fig. 5C, D**). The same trends also hold for the individual macroscopic parameters for the ON/OFF model; we observe a higher K_on_ but similar K_off_ at active locations regardless of the promoter identity (**Fig. S5A, B**).

In summary, we found that the regional chromatin environment and the local promoter identity independently control expression. While promoter identity has a large effect on expression mean, the activeness of the chromatin environment has a large effect on MIN.

## Discussion

Taken together, our results suggest that there is a strong, inherent relationship between mean mRNA levels and expression noise, but that this relationship can be modified by the epigenetic properties of different chromosomal environments. About sixty percent of all noise in mRNA expression is set solely by a gene’s mean level of mRNA production through a power law relationship, regardless of the identity or genomic location of the local CRSs. This observation suggests strong mechanistic constraints on noise that originate from fundamental properties of the transcriptional machinery. However, our data also show that the different properties of chromosomal environments can, in part, uncouple expression noise from mean levels of expression. We found that changing the dynamics of the transitions between ON/OFF states has an orthogonal effect on mean and mean-independent noise. This mechanism could allow natural selection to select for noisy expression without changing a gene’s mean expression level.

Our data also suggest that local and regional sources of cis-regulatory information act independently on a gene’s expression noise. Reporter genes have lower noise when integrated at active regions of the genome than at repressed regions, and this is true regardless of the identities of their promoters. Local and regional cis-regulatory information also control different aspects of transcriptional dynamics. The local cis-regulatory information controls the basic rates of transcription, regardless of its chromosomal environment, whereas the chromosomal environment has a large effect on the activation and inactivation dynamics of the gene, and consequently has a larger effect on mean-independent expression noise. We speculate that more active chromosomal environments have higher local concentrations of the components of the transcriptional machinery available, resulting in less heterogeneous transcription among individual cells. Meanwhile at more repressed genomic regions, the binding of the transcriptional machinery is a rarer event, increasing time intervals between transcriptional bursts and promoting heterogeneous expression in the population. This model suggests the possibility for predicting the noise in gene expression based on genomic location.

## Author Contribution

S.Z. and B.A.C. conceived the project. S.Z. performed experiments and analyzed the data. S.Z. and Z.P. created image acquisition pipelines. S.Z. and B.A.C. wrote the paper with input from all the authors.

## Acknowledgement

We are grateful to Sara Rouhanifard and Arjun Raj for providing us protocol and initial reagents for clampFISH experiments. We are also very grateful for Clarice KT Hong for providing the 4C data. We also thank the members of the Cohen Lab for their helpful comments and critical feedback on the manuscript; the Genome Engineering and iPSC Center for kindly allowing us to use their flow cytometer for cell sorting. This work was supported by NIH grants R01-GM092910 (to B.A.C).

## STAR Methods

### LEAD CONTACT

Further information and requests for resources and reagents should be directed to and will be fulfilled by the lead contact: Barak A Cohen.

### MATERIAL AVAILABILITY

The materials will be available upon request. DATA AND CODE AVAILABILITY

The file for the raw mRNA counts is provided in the Supplemental Table 3. The file for fitted rates is provided in the Supplemental 4. Raw images will be available upon request. The code will be stored on Github after the peer review process.

### KEY RESOURCES TABLE

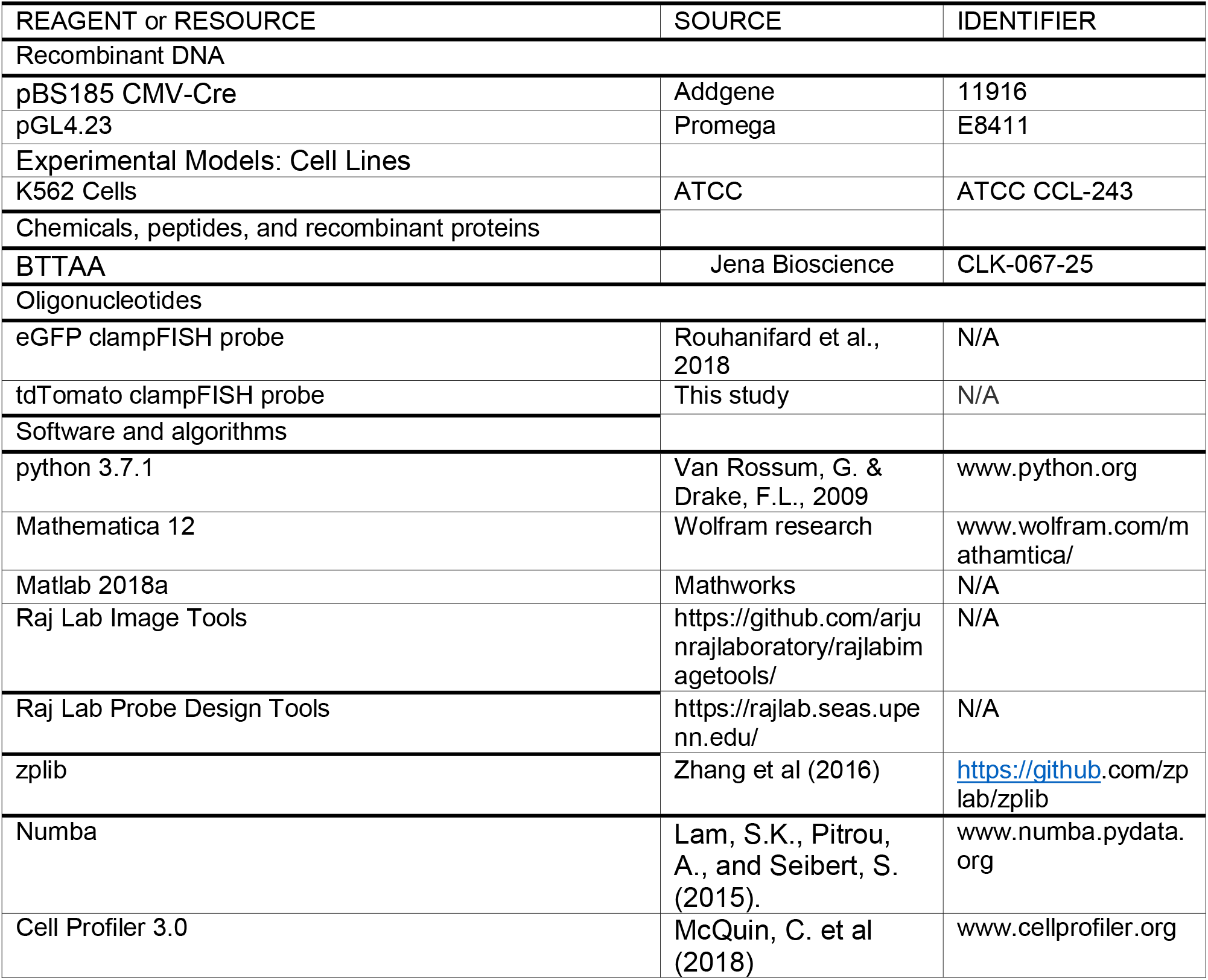

### EXPERIMENTAL MODELS AND SUBJECT DETAILS

#### Landing Pad Design and Cell Line Maintenance

K562 cell lines carrying landing pads constructions were taken from our previously published work (Maricque et al., 2018). There are several important features of the landing pad cassette: First, a pair of asymmetric Lox sites is in the cassette, allowing integration of different DNA sequences into the same location. Second, a 12-bp DNA sequence barcode was cloned downstream of one of the Lox sites, allowing the mapping of the genomic location of each landing pad. Those locations were mapped individually to the Human hg19 genome. Twenty-two different landing pads were chosen for the experiments presented in this work.

Landing pad K562 cells were cultured using a medium consisting of Iscove’s Modified Dulbecco’s Medium (IMDM) + 10% Fetal Bovine Serum (FBS) + 1% non-essential amino acids + 1% pen/strep.

#### Plasmid Design and Construction

The transfer vectors containing promoter cassette were subcloned with the following steps: First, the sequence of each promoter (PSMB2, CHMP2A, and HBZ) was extracted from the Ensembl genome browser (Yates et al., 2020). The 700 bp sequence before the TSS for each promoter was synthesized by Twist Bioscience. Second, the synthesized fragment was cloned into a pGL4.23 transfer vector backbone driving a tdTomato reporter using NEB HIFI assembly.

#### Cre-mediated promoter integration

Promoter swap was achieved using Cre-mediated recombination using the asymmetrical loxP, loxFAS sites in the landing pad. The transfer vector with different promoters (PSMB2, CHMP2A, HBZ) and the plasmid encoding Cre recombinase (pBS185 CMV-Cre, Addgene 11916) were transfected using Invitrogen Neon Transfection System (Life Technologies). For each transfection, we electroporated 4 ug of transfer vector containing different promoters with 4 ug of plasmid encoding Cre recombinase into 1.2 million K562 cells from a landing pad cell line. After transfection, we grew the cells for seven days, and we used FACS to sort single tdTomato-positive K562 cells into 96-well plates. Individual clones were expanded in culture medium for 1-21 days.

#### clampFISH Probe Design and Generation

ClampFISH probes for the reporter genes were designed using the Raj Lab Probe Design Tool (rajlab.penn.edu)(Rouhanifard et al., 2018). Each probe was broken into three arms to be synthesized by IDT. The 5’ of the left arm is labeled by a hexynyl group, and the 3’ of the right arm is labeled by NHS-azide. The right arm fragment was purified by HPLC. All three components were resuspended in nuclease-free H2O to a concentration of 400 uM. The three arms were ligated by T7 ligase (NEB, Cat# M0318L), at 25 C overnight. then purified using the Monarch PCR and DNA cleanup Kit (NEB, Cat# T1030S) and eluted with 40 ul of nuclease-free water. After the ligation, each probe is stored at −20 C. The list of oligos used in this paper can be found in Supplemental Table S2.

#### clampFISH Experimental Procedure

ClampFISH was performed according to the suspension cell line protocol of clampFISH (Rouhanifard et al., 2018). 1.2 million cells were collected and fixed in 2 mL of fixing buffer containing 4% formaldehyde for 10 min, then permeabilized in 70% EtOH at 4 C for 24 hours. The primary ClampFISH probes were then hybridized for 4 hours at 37 C in the hybridization buffer (10% Dextran Sulfate, 10% Formamide, 2X SSC, 0.25% Triton X). After hybridization, cells were spun down gently at 1000 rcf for 2 min. Cells were washed twice with the washing buffer (20% formamide, 2X SSC, 0.25% Triton X) for 30 min at 37 C. The secondary probes were then hybridized to cells at 37 C for 2 hours and the cells were then washed twice with washing buffer for 30 min at 37 C. The primary and secondary probes are “clamped” in place through a click reaction (CuSO4 75 uM, BTTAA 150 uM, Sodium Ascorbate 2.5 mM in 2X SSC) for 20 min at 37 C. The cells were then washed twice in the washing buffer at 37C for 30 min each wash. Then, the cells were hybridized with the hybridization buffer with tertiary probes for 2 hours at 37C. We complete 6 cycles of hybridization for all our experiments. After the final washes, cells were incubated at 37 C with 100mM DAPI for 20 min, washed twice with PBS, resuspend in the anti-fade buffer, spun onto a #1.5 coverslip (part number) using a Cytospin cytocentrifuge (Thermo Scientific), mounted onto a glass slide, sealed in antifade buffer with sealant, and stored at 4C.

#### Imaging

All images were taken within 72 hours of mounting of the slides. All images were captured by a 63X oil-immersion inverted wide field scope (Leica DMi8) with customized stage, camera (Andor Zyla 5.5) and filter sets (Chroma VCGR-SPX-P01). Automated image acquisition script was achieved through a custom imaging system developed in the previous publication (Zhang et al., 2016). We acquired z-stacks (1.2 μm between stacks) of stained cells.

### QUANTIFICATION AND STATISTICAL ANALYSIS

#### Image Processing

Once the images were collected, we perform a maximum z-projection to reduce the z-stacks into 2D images. We then used CellProfiler 3.0 for segmenting cells (Carpenter et al., 2006; McQuin et al., 2018). Briefly, trans images were first preprocessed to enhance the edges of the cells using the Prewitt algorithm. Then nuclei were identified with global minimum cross-entropy thresholding from DAPI images. Relying on the size and location of the nucleus, cell boundaries were segmented with a Watershed Algorithm. Once the cells were segmented, we then used the rajlabimagetools to quantify RNA FISH spots (https://github.com/arjunrajlaboratory/rajlabimagetools/). At this step we manually inspected each segmented cell and removed poorly segmented cells and adjusted the threshold for signal detection.

#### Gillespie Simulation for the ON/OFF Model

To investigate possible explanations for the observed differences in mean-independent noise associated between different chromosomal environments, we employed a two-state stochastic model for extrapolating the transcriptional dynamics. A parameter sweep of a stochastic model is computationally expensive but makes the least assumptions of the relationships among the kinetic parameters. To set up the simulation using the previously established framework, we first constructed the Chemical Master Equation (CME) for the ON/OFF model (Paulsson, 2005; Raj et al., 2006; Sherman and Cohen, 2014; Sherman et al., 2015), and by convention, we separate the Chemical Master Equation into two equation:

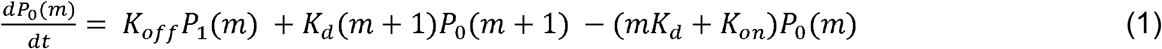

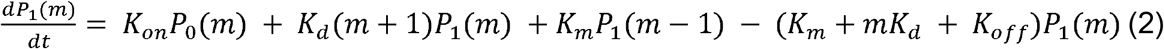

*P*_l_(*m*) and *P*_0_(*m*) denote the probabilities of having m molecules in the system in the ON and OFF states.

After establishing the CME for the ON/OFF model, we used the Gillespie Algorithm to simulate 8.8 million sets of parameters (Gillespie, 1976). We chose the estimated physiological range of the rates of activation, transcription, and mRNA degradation (K_on_: 0.00001 - 50 s^−1^, K_off_: 0.00001-50 s^−1^, K_m_: 0.002-1 s^−1^, K_d_:0.0002 - 0.003 s^−1^) (Chubb et al., 2006; Rodriguez et al., 2018; Suter et al., 2011). K_on_ and K_off_ were first broken down into intervals (0.00001-0.001 s^−1^ and 0.001-50 s^−1^). For the first interval, 37 values for each parameter were chosen logarithmically to ensure finer sampling for the smaller values. For the second interval, 50 values for each parameter were chosen logarithmically. Each set of parameters was simulated for 1×10^3^ individual trajectories till reaching stationary distribution, and the final distribution was recorded. The simulation was done on the Washington University High Throughput Computing Facility at Center for Genome Sciences (https://htcf.wustl.edu) with 200 cores with 2 Gb of RAM per core.

#### Fitting experimental data to simulated results

To fit our experimentally measured mRNA distribution with the ON/OFF model, Kolmogorov-Smirnov test was employed to determine the closest simulated distributions to the experimentally measured distribution. All sets of parameters that accept the null hypothesis for the K-S test (p > 1×10^−2^) are reported for subsequent analyses.

#### Epigenome data analysis

For the epigenetic data analyses at different integration sites, we considered the boundaries of interest as the 2500 bps flanking the integrated site. We then downloaded various K562 epigenome datasets (full list of sources in Table S5). For H3K9ac, H3K27ac, H3K9me3, H3K4me3, H3K27me3, DNase-seq we used pyranges (Stovner and Sætrom, 2020) module to overlay the signal with the integration sites. The ChromHMM and Segway combined segmentations for K562 cells were downloaded from UCSC genome browser and overlaid with the integrated sites.

#### TRIP data analysis

We downloaded TRIP data for mouse embryonic stem cells (mES cells) from Akhtar et al. (Akhtar et al., 2013) and the ChromHMM segmentation for mES cells from Pintachuda et al. (Pintacuda et al., 2017) (full list of sources in Table S5). We lifted the TRIP data from mm9 to mm10 using the UCSC Liftover Tool (Kent et al., 2002). We then overlaid the TRIP expression data with the ChromHMM segmentation using pyranges (Stovner and Sætrom, 2020).

### DATA AND SOFTWARE AVAILABILITY

The data for the raw mRNA counts is provided in the Supplemental Table 3. The data for fitted rates is provided in the Supplemental 4. Raw images will be available upon request. The code will be stored on Github after the peer review process.

---

**Figure S1.**
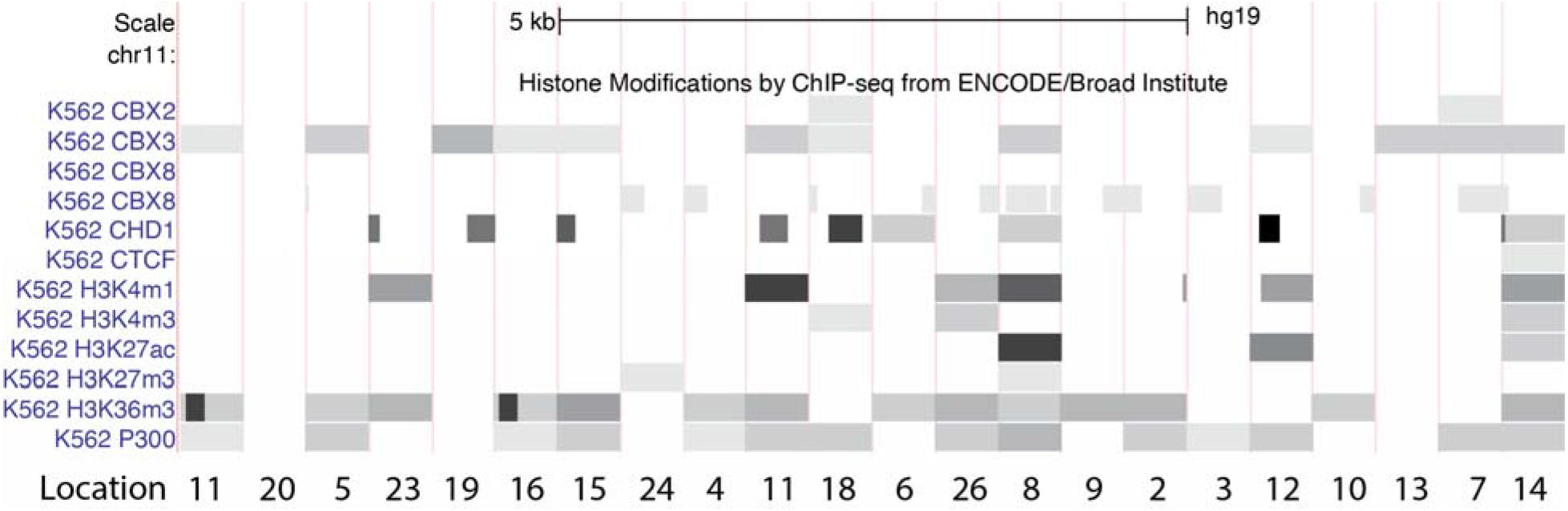
Landing pad locations have diverse epigenetic landscapes, related to Figure 1. UCSC genome browser view of the investigated locations with ChIP-seq data for several transcription factors and epigenetic modifications.

**Fig S2.**
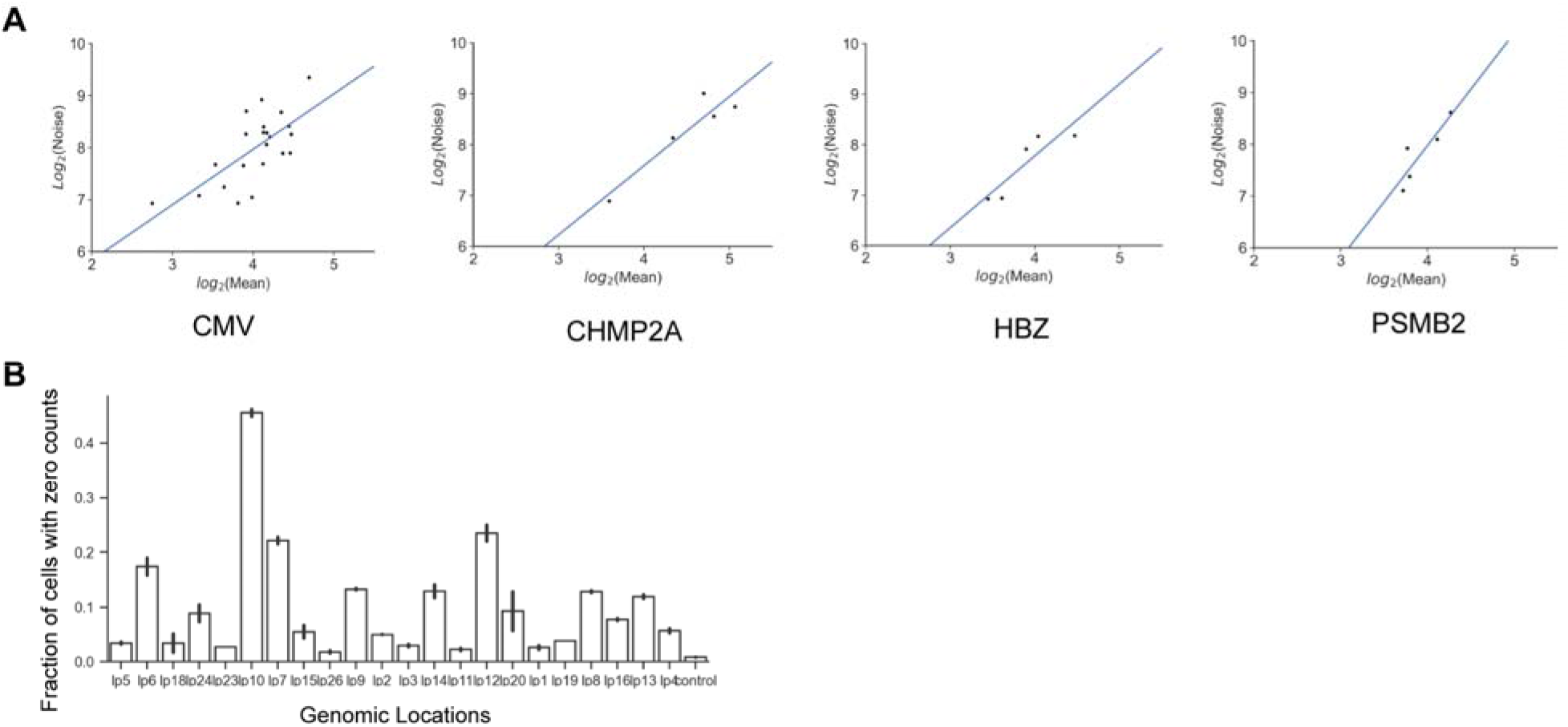
The Power-law relationship for different promoters at different genomic locations are largely the same and suggests a slow dynamics, related to Figure 1, 5. **(A)** Power-law relations for expression mean and noise for different promoters at different genomic locations in K562 cells. **(B)** The fraction of cells with no mRNA molecule labeled, compared to ACTB intron control. Error bar indicates the standard error from two biological replicates.

**Fig S3.**
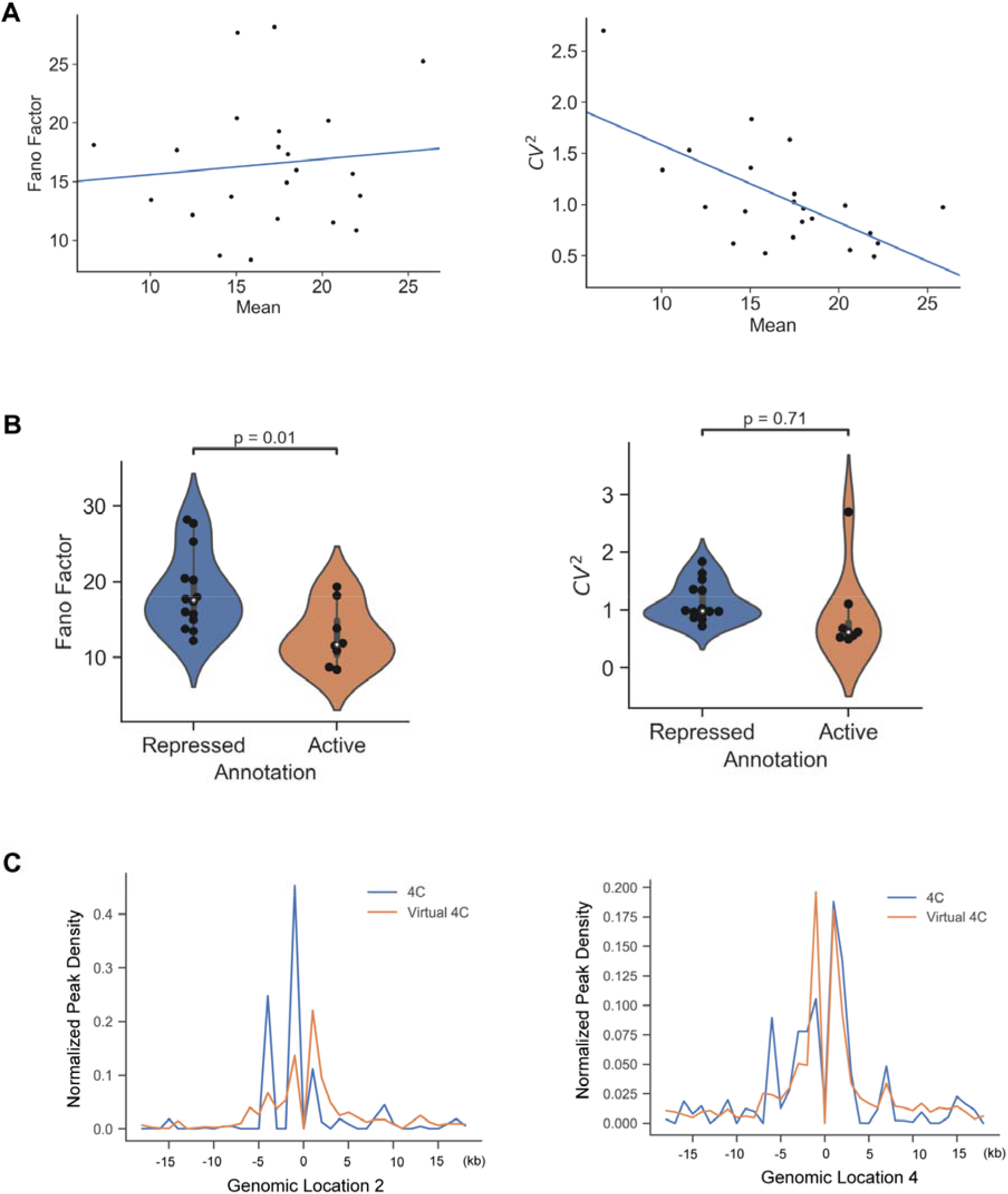
Specific noise metric does not change our observation, related to Figure 2. **(A)** Power-law relations for expression mean and Fano factor and CV^2^ for CMV promoters at different genomic locations in K562 cells. **(B)** Box plot showing Fano Factor and CV^2^ as noise metric for active and repressed locations. **(C)** Comparison of peak density between 4C (blue line) and Hi-C derived virtual 4C (orange line) for Location 2 and Location 4. Each peak represents the normalized read density centered on the insertion site.

**Fig S4.**
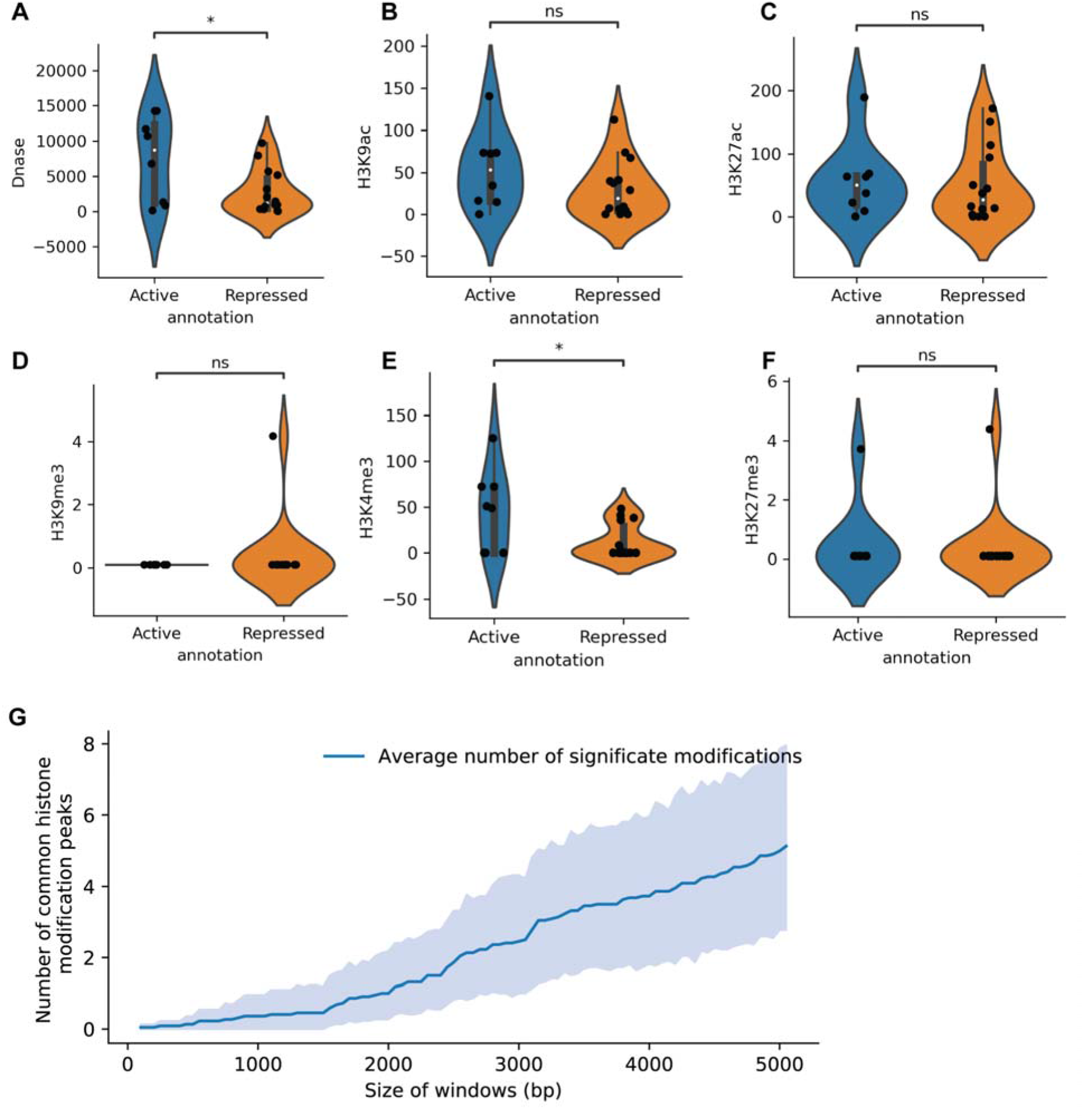
Epigenetic Analysis for different genomic locations, related to Figure 3,4. ChIP-seq datasets of histone modification are downloaded from the ENCODE and are correlated with active and repressed regions. **(A-C)**. Violin plot showing active histone modifications peaks 5kb surrounding the integration site at active and repressed locations. **(D-F).** Violin plot showing repressive histone modifications peaks 5kb surrounding the integration site at active and repressed locations. **(G)** Number of histone ChIP-seq peaks (H3K9ac, H3K27ac, H3K9me3, H3K4me3, H3K27me3) and DNase-seq peaks within a given window centered on the integration site for 22 genomic locations in K562 cells Solid blue line: average number of peaks, blue shade: 95% C.I.).

**Fig S5.**
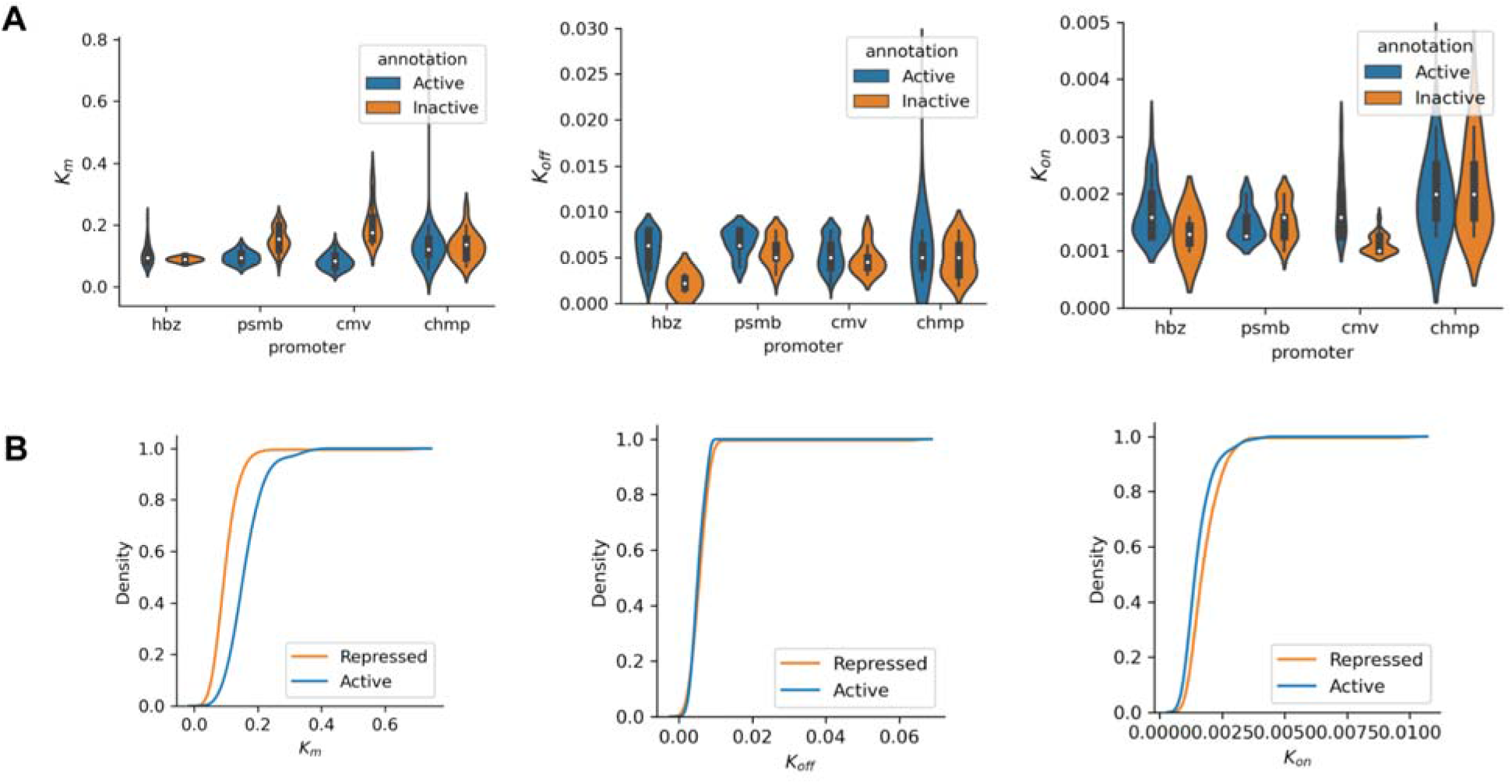
Fitted rates for different promoters and different genomic locations, related to Figure 5. **(A)** Different rates (K_m_, K_off_, and K_on_) of the ON/OFF model for different promoters at active and repressed locations. **(B)** Cumulative distribution function for different rates at active and repressed locations

